# Modulation of tactile sensitivity in the lower limbs during goal-directed movements

**DOI:** 10.1101/2025.07.28.667135

**Authors:** Fabian Dominik Wachsmann, Katja Fiehler, Dimitris Voudouris

**Affiliations:** Experimental Psychology Justus Liebig University Giessen Otto-Behaghel-Str. 10F, 35394, Giessen, Germany

**Keywords:** Tactile Suppression, Tactile Masking, Sensory Integration, Postural Control, Kicking

## Abstract

Tactile sensitivity drops in a moving than static limb due to a combination of central, predictive mechanisms and peripheral effects. This suppression is dynamically modulated during movement, as shown during upper-limb actions, yet little is known about its implication during complex lower-limb movements. We investigated tactile sensitivity during naturalistic kicking by delivering vibrotactile probe stimuli to the balancing and kicking feet at different movement phases. In Experiment 1, participants kicked a suspended ball while tactile sensitivity was probed at movement *onset*, mid-*swing*, ball *contact*, and *after*-contact. Results revealed distinct modulation patterns in each foot. When transitioning from bipedal to unipedal stance, tactile processing at the balancing foot was particularly suppressed but at the kicking foot it improved, suggesting concurrent modulation across the two legs depending on their motor function. Tactile sensitivity remained rather invariant at other time points, but was strongly suppressed on the kicking foot at the moment of ball contact. The strength of this suppression correlated with kicking speed, which could reflect either stronger predictive control or stronger peripheral processes that mask the vibrotactile probe. To test these, a new set of participants held their foot still while a ball collided with it at high or low speed. Suppression was greater with faster ball contacts, revealing that peripheral processes can modulate tactile processing. These findings show that lower-limb tactile sensitivity during goal-directed leg movements can be concurrently modulated across the legs, presumably reflecting an interplay between central sensorimotor processes guiding the movement and peripheral processes affecting sensitivity.

**Significance Statement:** Tactile sensitivity is known to fluctuate during movement, but little is understood about how it is tuned during complex lower-limb actions. Using a naturalistic ball-kicking task, we reveal distinct modulation patterns in the balancing and kicking feet, showing that postural and guiding demands dynamically shape tactile sensitivity. We further demonstrate that the strength of tactile modulation is influenced by peripheral processes, such as tactile masking. These findings highlight that lower-limb tactile processing can be flexibly and concurrently modulated in the two legs during state transitions that impose different sensorimotor demands for complex natural behavior.

## Introduction

Tactile sensitivity is typically reduced on a moving limb compared to when the same limb is resting. This suppression stems primarily from predictive mechanisms, in which motor commands anticipate future sensory states, leading to suppression of associated sensory feedback (Blakemore et al., 2000). This predictive process explains why self-produced sensations, like tickling (Blakemore et al., 2000) or the strength of self-applied forces (Shergill et al., 2003), feel weaker during active tasks. However, externally-generated sensations, which cannot be predicted based on motor commands, are also suppressed on a moving limb (Arikan et al., 2024; Fuehrer et al., 2022). This led to the idea that the suppression of externally-generated sensations arises from a general gating mechanism unrelated to prediction (e.g., Kilteni & Ehrsson, 2022). However, accumulation of findings demonstrates that suppression of externally-generated sensations reflects the integration of feedforward predictions with online sensory feedback (Fuehrer et al., 2022; Mouchnino et al., 2015). For example, tactile suppression during grasping is reduced when tactile input is important for object manipulation (Voudouris et al., 2019; Voudouris & Fiehler, 2022). Suppression is also temporally modulated during goal-directed arm movements (Colino & Binsted, 2016; Fraser & Fiehler, 2018), with recovered sensitivity when somatosensory feedback becomes critical (Voudouris & Fiehler, 2021). Tactile sensitivity can also change in the leg during whole-body actions, such as walking (Pearcey & Zehr, 2019). For instance, it can improve when preparing to avoid a visual perturbation (Wachsmann et al., 2025a) or just before initiating a step (Mouchnino et al., 2015), reflecting upregulation of task-relevant tactile input for postural stability. Overall, these results demonstrate the flexible, context-dependent modulation of tactile sensitivity during movement.

An important distinction between upper-limb and whole-body actions is that the latter require the retention of upright stance. This on its own requires a cascade of sensorimotor processes from multiple effectors to ensure a stable posture and the successful execution of the overarching body action. For instance, when retaining bipedal posture, the sensorimotor system should control sensory processing from two separate effectors (legs) that may sometimes even be exposed to different dynamics. One such example occurs during postural transitions, like walking or kicking, when the two effectors need to simultaneously coordinate the redundant degrees of freedom based on the available sensory information and feedforward predictions, to ensuring stable upright stance and successful performance.

Considering these, we are here interested in the regulation of tactile processing from both legs during postural transitions. We focus on ball kicking, which involves complex sensorimotor processes for both feet. The balancing foot supports posture, while the kicking foot executes a goal-directed movement. Performance is associated with balancing ability (Chew-Bullock et al., 2012) and a tradeoff between speed and accuracy of the kicking leg (Kellis & Katis, 2007). Meanwhile, cutaneous and other sensory signals inform about postural disturbances (Peterka, 2002). We examined tactile processing in the balancing and the kicking foot during postural transitions, when each leg serves different functional purposes to foster successful behavior. We propose two main hypotheses. First, for the balancing foot, we expected reduced tactile sensitivity during the *swing* phase of the kicking foot compared to other moments because the transition from a stable bipedal to an unstable unipedal stance can reduce tactile sensitivity (Wachsmann et al., 2025b). Second, for the kicking foot, we predicted enhanced tactile sensitivity during the *swing* phase that would reflect uptake of somatosensory input to facilitate the guidance of the moving foot to the ball (Voudouris & Fiehler, 2021). We further hypothesized reduced sensitivity in the kicking leg at ball contact due to masking (Fraser & Fiehler, 2018). To examine this deeper, we tested whether the speed of a moving ball can modulate tactile sensitivity on a static leg at various times around ball-foot contact. Based on earlier research (Abramsky et al., 1971; Kirman, 1984), faster collisions should cause stronger suppression due to peripheral processes masking the sensations in a limited window around contact.

## Methods Experiment 1

### Participants and apparatus

We recruited 23 young (25.83 ± 3.69 years old, range: 21-36; height: 175.52 ± 10.23 cm; 8♀, 15**♂**) healthy participants. Participants were free from any known neurological or musculoskeletal issues at the moment of the experiment and had normal or corrected-to-normal vision. Upon their arrival at the lab, they provided their signed informed consent. At the end of the experiment, participants received either 8€/hour or course credits. The study was approved by the local ethics committee of the Justus Liebig University Giessen and was conducted in accordance with the “World Medical Association Declaration of Helsinki” (except for §35, pre-registration) (“World Medical Association Declaration of Helsinki”, 2013).

Two custom-made vibrotactile stimulation devices (Engineer Acoustics Inc., Florida, USA), referred to as “tactors,” were attached to the glabrous skin between the third and fourth metatarsal heads of both feet —an area known for its sensitivity to vibrotactile stimuli (Hennig & Sterzing, 2009). The tactors comprised a small housing (22 x 45 x 5 mm) and a round actuator (6 mm diameter) that can generate vibrotactile stimuli of fine-grained amplitude ranging from 0.00316 – 0.632 mm in steps of 0.00316 mm (Figure 1a). We used these tactors to generate brief tactile stimulations that served as a proxy to examine tactile processing from the stimulation site. This is based on findings and procedures from several previous studies that examined tactile sensitivity during upper-limb (Buckingham et al., 2010; Fuehrer et al., 2022; Voudouris & Fiehler, 2017, 2021) and lower body movements (Menz et al., 2006; Wachsmann et al., 2025a, 2025b). Care was taken to ensure that loose wires or clothing did not interfere with the perception of vibration or hinder task performance.

**Figure 1:**
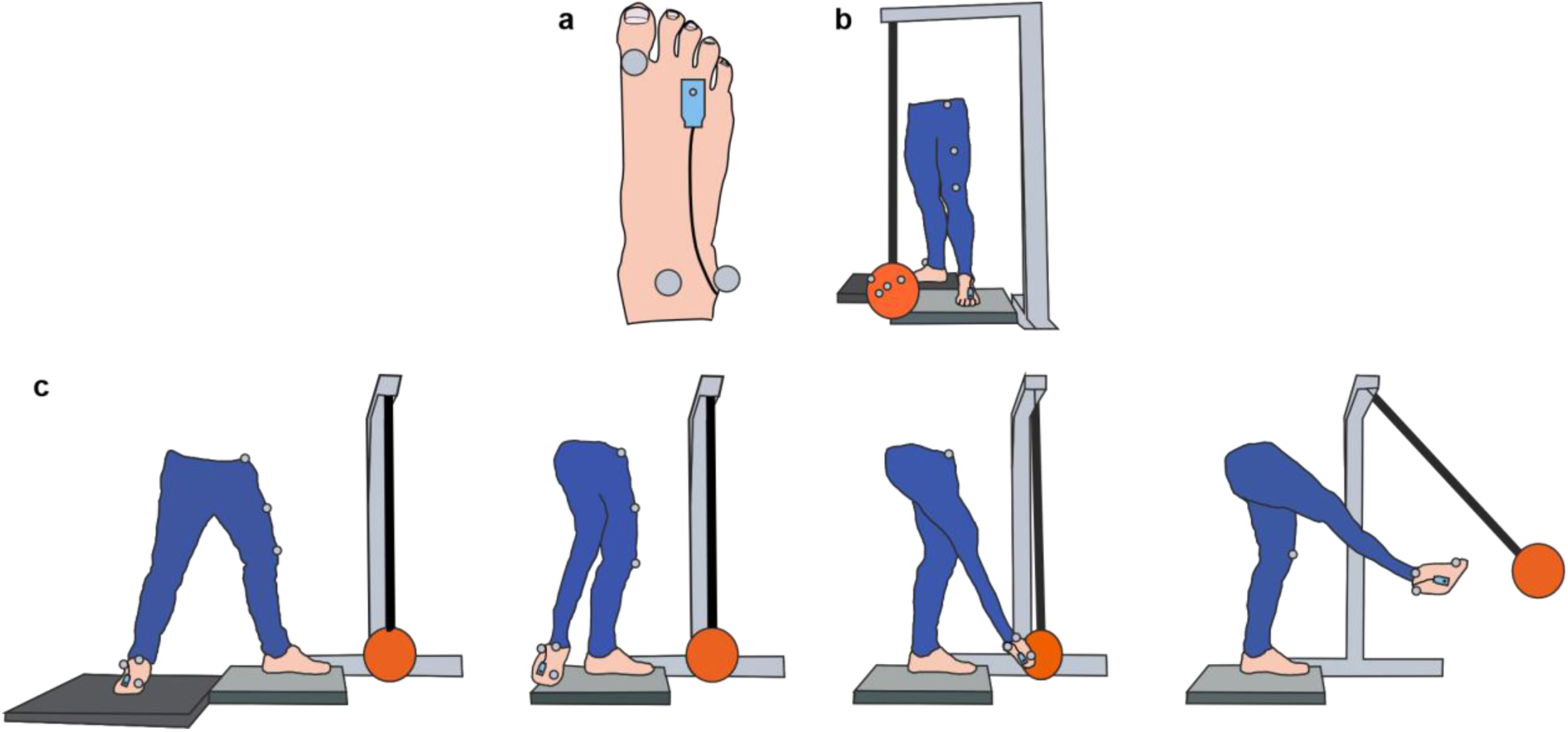
Setup Experiment 1. (a) Top view of the right foot with the tactor placed at the metatarsophalangeal joint between the 3^rd^ and 4^th^ toes. (b) Experimental setup, with the ball suspended from a beam with the 4 markers attached to it and a platform where participants stood on. The three markers, as well as the tactile device, are depicted for the left leg. (c) Sketch of the starting position, mid-swing, ball contact, and after-contact time points (from left to right).

During the experiment, participants placed their left foot on a platform and kept it there. The position of each leg was recorded with reflective markers (spheres of 10 mm diameter) that were placed on certain anatomical landmarks of each leg. Specifically, for the kicking leg, the markers were placed at the malleolar fork, the lateral malleolus, and the first metatarsophalangeal joint (Figure 1a). For the balancing leg, the markers were attached at the knee cap, the *spina iliaca anterior superior,* and at the middle of the line connecting the previous 2 markers (Figure 1b). In addition, four reflective markers were placed on a foam ball (21 cm diameter, 290 g), which was suspended from a beam in front of the participant. The top part of the beam was positioned 115 cm above standing level, and the ball was suspended 1-2 cm above the standing level to enable free kicking. The variance in suspension level was due to the rope sometimes winding up and thus slightly shortening during several kicks. All ten markers were tracked at 300 Hz using eight Vicon Vero 2.2 (Vicon Motion Systems, Oxford, UK) cameras, with data captured and processed using Vicon Nexus 2.15 software. The marker data were streamed into MATLAB (version 2023b, The MathWorks Inc., Natick, Massachusetts, USA) via the Vicon DataStream SDK (version 1.12.0). A schematic depiction of the setup, including the positioning of the tactors and markers can be seen in Figure 1a-b.

### Procedure

The experiment consisted of four main blocks: two baseline blocks to assess tactile sensitivity in each foot during rest, and two experimental blocks to assess changes in tactile sensitivity during kicking. Before these four blocks, participants underwent a familiarization phase during which they got accustomed with the tactile stimuli and practiced the kicking trials.

During the familiarization phase, participants stood on the platform with their feet at shoulder-width apart. They performed five trials, each of which included a vibrotactile stimulus (50 ms, 250 Hz) at either the balancing (always left) or the kicking (always right) foot. The stimulus intensity ranged from weak (peak-to-peak amplitude: 0.00316 mm) to strong (peak-to-peak amplitude: 0.284 mm), allowing participants to become familiar with the sensations. All participants were able to detect at least one of these stimuli. After these five trials, participants held their kicking foot against the hanging ball, which allowed us to determine the minimum distance between the foot and the ball markers. This was important to later define the moment when the kicking foot contacted the ball (see below). Specifically, participants had to touch the ball with the medial side of their right foot, as if they were performing a kick. As soon as this was visually confirmed by the experimenter, the positional data from the markers on the foot and the ball were collected for 1 second. We then determined the median distance between the averaged positions of the right foot and of the ball markers over the recording. Participants then completed six practice kicking trials. They were instructed to take a kicking-ready stance, with the balancing foot in front and the kicking foot positioned further behind on another platform of similar height (Figure 1c). Tape markings were placed on the platforms to ensure consistent foot positioning between trials for each participant. In each trial, participants were asked to kick the suspended ball and attempt to replicate the same movement across all kicks. During the six practice kicking trials, participants also received high-intensity vibrations (0.632 mm) at three different time points: when lifting the kicking foot toward the ball, at ball contact, and 250 ms after ball contact. The first three trials involved vibrations at the balancing foot, and the next three trials included stimuli at the kicking foot. Each trial was initiated by the experimenter pressing a button followed by a brief tone after one second, indicating that the participant should execute the kick. The kick had to be completed within 15 seconds from the onset of the tone. The onset of movement was defined as the first moment when the average position of the markers on the kicking foot moved more than 1.5 cm from their starting position, which was defined as the average position of the foot markers during the 100 ms preceding the tone. Ball contact was detected when the Euclidean distance between the kicking foot and the ball was within 1.5 cm of the previously determined minimum foot-ball distance. This familiarization procedure lasted approximately 5 minutes.

After these familiarization trials, participants were randomly assigned to one of four possible sequences of blocks, each block assessing tactile sensitivity on one of the feet under a baseline and a kicking session. The order of the four blocks was randomized with the constraints that (a) the baseline blocks had to be presented sequentially either as the first or the last pair of blocks, and (b) the order of the tested feet was identical between baseline and kicking blocks. The four sequences were balanced across participants. Baseline measures were taken in sitting with both feet touching the ground. In each baseline trial, a tactile stimulus was delivered to the foot, and the experimenter asked the participants to verbally report whether they felt a vibration or not. The experimenter then logged the response to the host PC, and after a random delay between 2 and 3.5 seconds, a new tactile stimulation occurred, followed by the experimenter’s question. The variable delay was introduced to minimize the temporal predictability of the stimulus. In each kicking trial participants had to first adopt the starting position. The experimenter then pressed a button that triggered an auditory go-cue 1 second later. This instructed participants to perform the kicking movement with their right foot within the next 15 seconds. A tactile stimulation occurred at one of four possible time points, which were fully randomized within each block. Three of these time points were identical to those used in the familiarization procedure: movement *onset*, ball *contact*, *after*-contact. The fourth time point occurred halfway through the *swing* phase of the kick (Figure 1c). This time point was estimated for each trial by calculating the swing movement time as the duration between movement onset and ball contact. We then calculated the average duration of the last five trials and used this as an estimate of the swing duration in the upcoming trial (e.g. Gertz et al., 2017; Arikan et al., 2021; Voudouris & Fiehler, 2022). This method provided robust estimates of movement times, resulting in a mean difference of 21 ± 21 ms between the intended and actual stimulation time for trials with stimulations planned during the swing phase (see Appendix 2 for a histogram). For the first five trials of each experimental block, we estimated the movement time based on the last trials of the practice kicks that were performed during the familiarization phase. For each of the two baseline conditions and for each of the four time points during the movement trials we presented 25 stimuli, totalling 250 trials per participant. This took about 45 minutes to complete. Participants could take breaks at any time during the experiment, but no one did.

The stimulus intensity for each trial was determined using a QUEST algorithm (Psychtoolbox Version 3.0.18; separate for each condition), which employs a Bayesian approach to estimate psychometric function parameters based on prior knowledge. This was done separately for each of the four blocks and separately for each participant. If suggested values were larger or smaller than those that the tactor could generate, we used the maximum (0.284 mm) or minimal (0.00316 mm) possible values, respectively. A Weibull function with β = 3.5 was used, as recommended for two-alternative forced choice (2AFC) tasks (Watson & Pelli, 1983). Prior mean and standard deviation estimates were based on a pilot study with young participants (N = 16), who were exposed to 50 stimuli of varying intensities while standing with their feet together. Lapse and guess rates were informed by another study on balance control during standing (Wachsmann et al., 2025b).

### Data analysis

We determined the detection threshold for each condition and participant as the intensity of the tactile stimulus that was determined by the QUEST after the participant’s response in the last trial of the respective block. As mentioned above, and if necessary, this intensity was corrected to fit within the possible stimulation range of the tactor. We were primarily interested in assessing how tactile sensitivity on either foot is modulated during a kicking action. Therefore, to account for individual differences in tactile sensitivity, we normalized each participant’s detection thresholds of the kicking and the balancing foot during kicking blocks to the respective thresholds obtained during the baseline blocks. This resulted in four Δthreshold values, with higher values indicating higher thresholds during kicking than baseline trials (i.e. suppression).

If a resulting Δthreshold value in a single condition was lying outside the median ± 3 * interquartile range, the participant was excluded from the analysis using that parameter. Each of the 8 normalized conditions was checked individually. This resulted in excluding three Δthreshold values, each from a separate participant, from the following conditions: kicking foot *onset*, kicking foot *swing*, and kicking foot *after*-contact. Please note that if a Δthreshold is excluded, the whole participant could not be used for the ANOVA (see below), which would require that value. This resulted in three participants being excluded.

To visualize the leg kinematics, we first smoothed the positional data with a 33 ms (10 frames) moving average window to reduce high-frequency artefacts. Then we determined the speed of each leg separately by numerical differentiating the average positional data of the markers on that leg. We did so separately per trial and condition, and we then averaged across those trials to obtain two average speed profiles (one per leg) for each participant. We then averaged across the participants, and we present these averages starting from 1 second before the onset of the right foot’s movement and until 1.5 seconds later. This is a time window that covers the preparatory and execution phases of the kick movement. In addition, we provide a time-normalized speed profile for each leg to further highlight the movement dynamics of each leg at the critical three time points of stimulation that occurred during the movement. To obtain the time-normalized speed profiles of each leg, we normalized the speed profile of every trial via linear interpolation to obtain a time course of 100 equal steps per trial, and then averaged across the 200 kicking trials of each participant.

### Statistical analysis

Our main interest is whether tactile sensitivity on the balancing and kicking feet is temporally modulated during a kicking action of the right foot. To assess this, we conducted 2 separate one-way repeated measure ANOVAs, one per foot, with 4 levels (time points of stimulation). For the statistical analysis we used the Δthresholds. Main effects were explored with post hoc comparisons using paired t-tests (two-sided), corrected using the Holm procedure.

## Results Experiment 1

### Psychophysics

Baseline tactile sensitivity on the two feet is depicted in Figure 2a. As can be seen, detection thresholds appear somewhat lower in the right (kicking) than left (standing) leg, but this difference is not systematic, while the variability across participants is considerable, yet similar between feet. This is in line with previous findings that indicate substantial individual differences in tactile sensitivity (Kozłowska, 1998). Therefore, examining the relative change of the detection thresholds between kicking and baseline trials allows us to reduce the impact of these individual differences and focus on the temporal modulation of tactile sensitivity.

**Figure 2.**
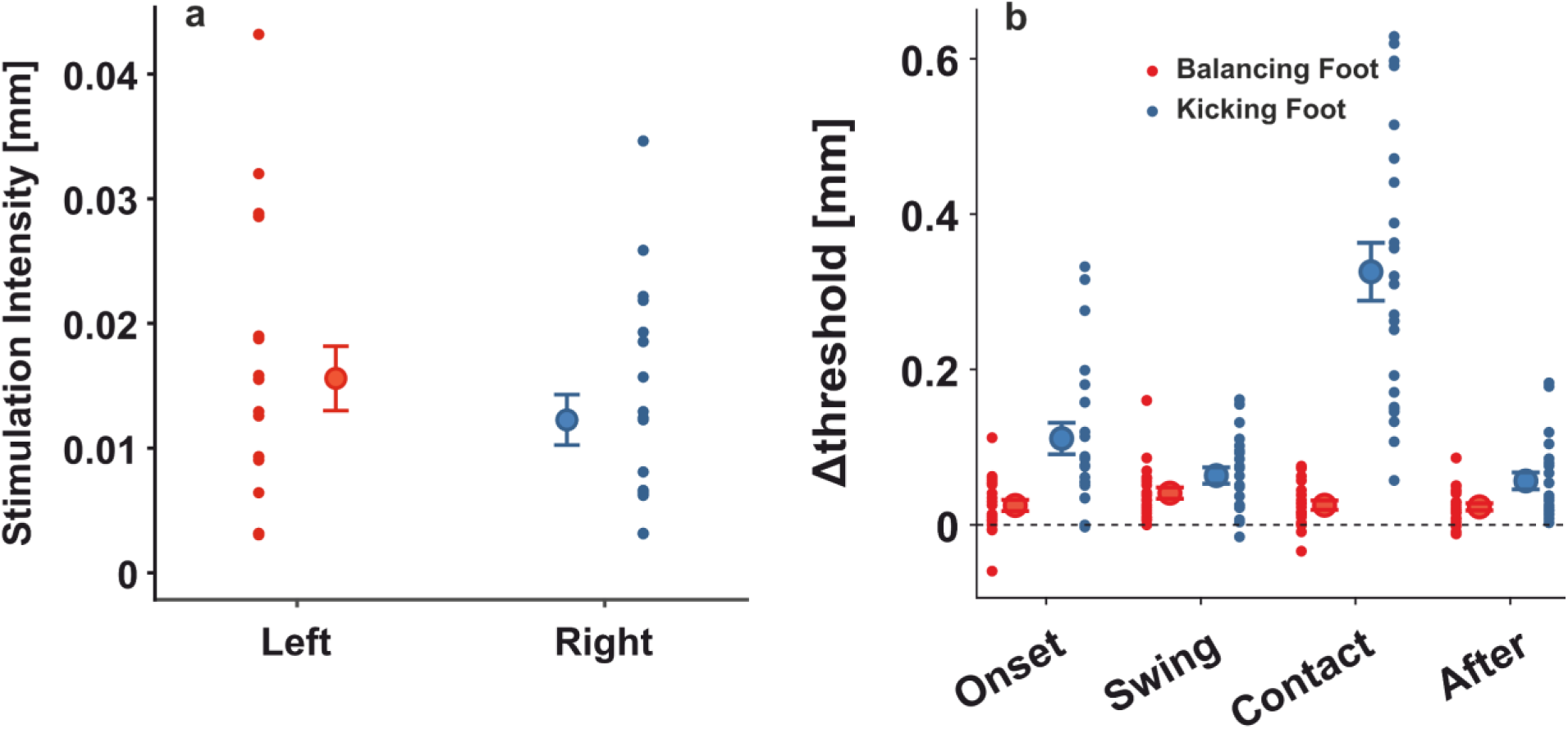
Temporal modulation of tactile sensitivity. (a) Raw detection thresholds obtained for the left (red) and the right foot (blue) during rest. Note that the left and right feet are always the balancing and kicking feet, respectively, during the experimental blocks. (b) Normalized detection thresholds as function of the four stimulation time points. The dashed line indicates baseline level thresholds. For both panels, red data corresponds to the balancing and blue to the kicking foot. Single dots represent single subject thresholds, while circles with error bars represent the mean and standard error of each condition, respectively.

During kicking trials, tactile sensitivity was suppressed on both feet (Figure 2b), as all Δthreshold values were larger than zero (all *t* >= 3.479, all *p* <= 0.002, all *η^2^* >= 0.116). This is in line with previous findings showing that standing itself can lead to reduced tactile sensitivity (Wachsmann et al., 2025b). In addition, and most importantly, tactile sensitivity was temporally modulated both on the balancing (*F*_3, 66_ = 3.604, *p* = 0.018 *η^2^* = 0.141) and the kicking foot (*F*_3, 60_ = 50.435, *p* < 0.001 *η^2^* = 0.716). For the balancing foot, sensitivity was poorest during the *swing* phase compared to all other 3 time points (*t* >= 2.524, *p* <= 0.049, *η^2^* >= 0.063), but none of the other comparisons was systematic (*t* <= 0.411, *p* = 1, *η^2^*<= 0.001). For the kicking foot, we found clear differences between all time points of stimulation (*t* >= 2.663, *p* <= 0.015, *η^2^* >= 0.042). Specifically, tactile sensitivity increased during the *swing* phase, before declining at the moment of contact and recovering again after the end of the kick. It is remarkable that tactile sensitivity decreased on the balancing foot but increased on the kicking foot during the *swing* phase, suggesting a concurrent modulation of tactile processing at the two feet, presumably reflecting the differential sensory processing to complete the two separate functional tasks that each leg has to solve for successful behavior.

### Kinematics

Figure 3a illustrates the average foot speed across all trials and participants, with the moment of movement *onset* and ball *contact* being highlighted with the two vertical purple lines. Thus, the swing phase is the period between these two lines. Figure 3b presents the time-normalized speed of each foot during *swing* phase averaged across participants, thereby accounting for variability in swing durations due to different kicking speeds. Individual normalized graphs for each participant are available in Appendix 1.

**Figure 3:**
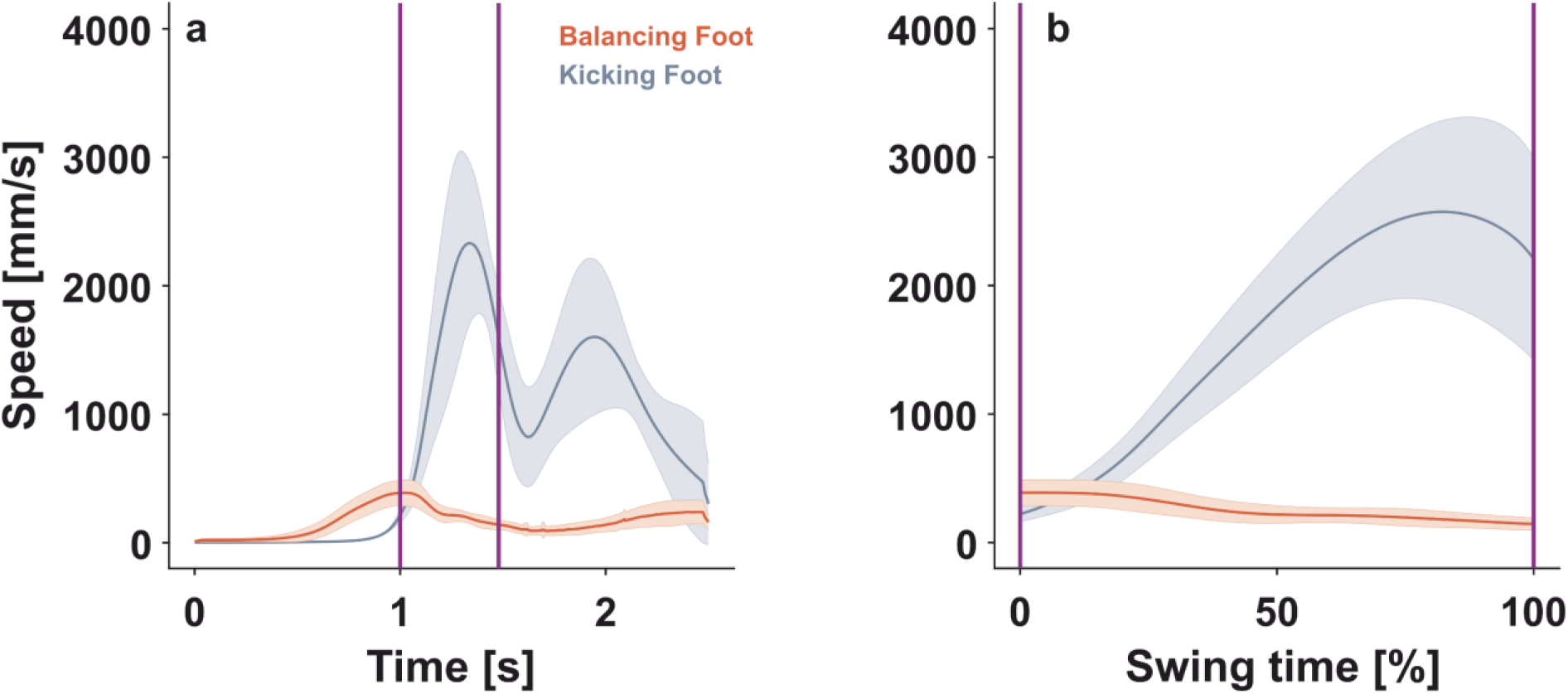
Kinematic results. (a) Time-course of the balancing (red) and kicking (blue) foot speeds from 1 second before movement onset until 1.5 seconds after movement onset. (b) Time normalized time-course of both feet speed during the swinging phase. Time-courses are averages across participants, with shaded bands indicating standard errors across participants. The first and second purple vertical lines indicate the moments of movement onset and ball contact, respectively.

As can be seen in Figure 3a, the action starts with the slow movement of the balancing leg, presumably indicating the postural adjustments that will enable the movement of the right leg. The speed of the kicking leg increased sharply ca. 500 ms later, reached its peak toward the end of the swing phase, and suddenly decreased its speed just before ball contact. This decrease in the movement speed appears consistent with patterns typically seen in goal-directed movements like reaching (Elliott et al., 2017). The speed of the kicking leg at the moment of contact ranged from 2157 mm/s (25th percentile) to 2579 mm/s (75th percentile), with a median of 2174 mm/s.

### Discussion Experiment 1

We examined tactile sensitivity on the two feet during a naturalistic ball kicking movement. We probed tactile sensitivity on both the balancing and kicking feet at four different time points of the action. First, we could replicate a previous finding that standing and the preparation of movement reduce tactile sensitivity on both feet compared to when simply sitting (Wachsmann et al., 2025a). For the balancing foot, we hypothesized that tactile sensitivity would decrease during the *swing* phase of the kick relative to all other moments of the action, based on previous findings showing that a rapid rise in balancing demands can reduce tactile sensitivity (Wachsmann et al., 2025b). Our findings confirm this hypothesis, providing further evidence for the idea that increased balancing demands when shifting from the bipedal to unipedal stance can compromise tactile sensitivity. This is at odds, though, with previous work showing improved tactile sensitivity in the leg shortly before a postural response to secure body balance, either when avoiding a visual perturbation (Wachsmann et al., 2025a) or when shifting weight to the balancing leg before initiating a step (Mouchnino et al., 2015). Therefore, the actual role of balancing demands on tactile sensitivity needs further investigation.

For the kicking foot, we hypothesized that tactile sensitivity would increase during the *swing* phase, driven by the need to guide the moving foot to accurately hit the ball. This is based on previous work from upper limb goal-directed action that showed improved tactile sensitivity on a reaching arm at moments when sensory guidance of the movement was more important (e.g., Voudouris & Fiehler, 2021; 2022; Fraser & Fiehler, 2018). However, it is also possible that tactile sensitivity would be reduced during the *swing* phase, because the high movement speed of the stimulated limb might lead to tactile suppression (Cybulska-Klosowicz et al., 2011). Our results do not lend evidence for this latter possibility, as sensitivity on the kicking foot was highest during the *swing* phase. This suggests that the possible effects of movement speed on tactile suppression may be less consistent. A striking finding of this experiment is that the detection thresholds in the kicking foot at ball *contact* were both highly elevated and rather variable across participants (see Figure 2b). These might arise from differences in kicking performance, particularly the variability in kicking speeds we observed in our data. A correlational analysis between kicking foot speed at the moment of ball *contact* and the strength of suppression (Δthreshold) from trials where the kicking foot was probed at that moment revealed a significant positive relationship (r = 0.467, p = 0.025, Appendix 3). We assume that tactile sensitivity is less influenced by the movement speed per se but rather by the difference in impulse caused by the collision with the ball, possibly through a masking mechanism that hinders the detection of the brief probe stimulus, similar to what has been suggested before (Abramsky et al., 1971; Kirman, 1984). To further explore the relationship between the speed of the kicking movement (masking stimulus intensity) and the resulting detection thresholds at around the moment of ball *contact*, we conducted a second experiment.

### Experiment 2

The reduction in tactile sensitivity at the kicking foot at the moment of ball *contact* could arise from the phenomenon of tactile masking. This phenomenon occurs when the perception of a tactile stimulus is masked by another stimulus that is presented in close spatial and temporal proximity (Fraser & Fiehler, 2018). Tactile masking can be categorized into three types based on their temporal properties. Forward masking occurs when the masking stimulus precedes the probe stimulus, typically within a window of 70 ms. Simultaneous masking happens when the masking and probe stimuli temporally overlap. Backward masking occurs when the masking stimulus follows the probe stimulus, typically within a window of 100 ms. The strength of forward and backward masking typically decreases with longer stimulus onset asynchrony (Abramsky et al., 1971; Kirman, 1984). In general, the strength of the masking effect is more pronounced when the masking stimulus is more intense (Craig, 1974; Kirman, 1984; Schmid, 1961). In Experiment 2 we explored whether the strong and variable suppression at the moment of ball *contact* found in Experiment 1 is driven by such masking effects irrelevant of any predictive mechanism (Chapman et al., 1987; Fuehrer et al., 2022) or sensory demands (Voudouris & Fiehler, 2021; Wachsmann et al., 2025a) that might be involved. We further tested whether tactile sensitivity at time points shortly before or after the moment of ball *contact* might have been influenced by backward- or forward-masking. To this end, a ball collided in different speeds with the participant’s static foot, and we assessed whether tactile sensitivity on that foot is influenced by the ball’s speed.

## Methods Experiment 2

### Participants and apparatus

We recruited again 23 young (22.04 ± 1.77 years old, range: 19-27; height: 176.09 ± 11.86 cm; 13♀, 10**♂**) healthy participants. None of them had participated in Experiment 1. The inclusion criteria, apparatus, procedure and analysis are identical to those of Experiment 1, except the details mentioned below. This time we had only one tactor at the right (receiving) foot to examine the masking effect of the collision. We also equipped the beam with 3 additional markers (one over the hanging for the ball, and two along the beam) to calculate the angle between beam and the ball. This was necessary to adjust the collision impulse. In addition, we built a cardboard blind to block participants’ view of the ball and of the experimenter. This measure was taken to ensure that no predictions about the speed of the ball and the timing of its contact with the foot could be established based on visible ball kinematics. A schematic depiction of the setup is provided in Figure 4.

**Figure 4.**
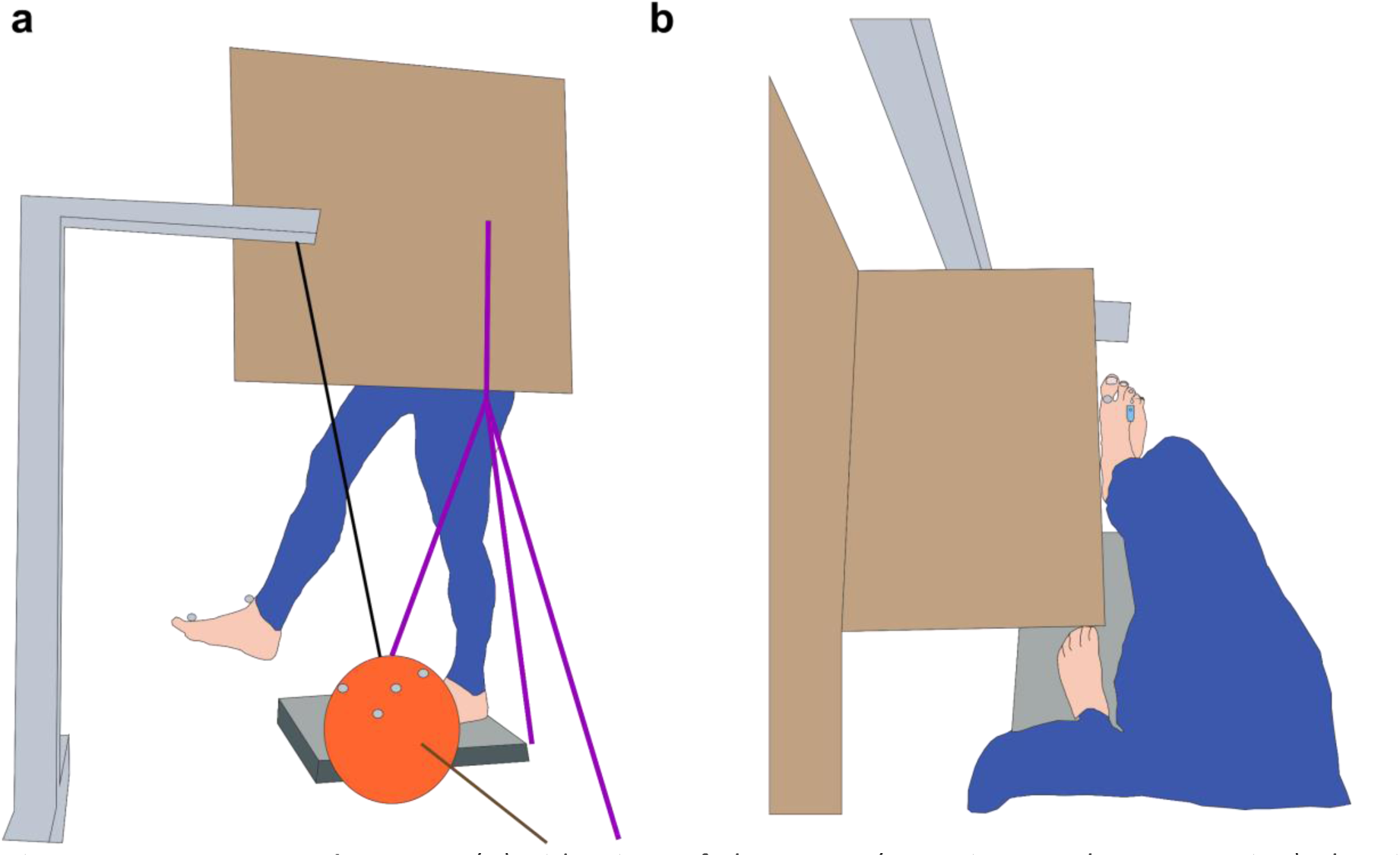
Setup Experiment 2. (a) Side view of the setup (experimenter’s perspective) showing the beam with the suspended ball, the purple tripod with the occluder attached, a participant standing in the experimental posture, and the four markers on the ball as well as two foot markers. (b) Top view (participant’ s view) with the occluder covering the ball and its possible trajectory during the experiment. Only one marker and the tactor are visible from this angle.

### Procedure

This experiment comprised four blocks of trials: Sitting baseline, standing baseline, high speed, and low speed. The familiarization phase as well as the determination of the minimal foot-ball distance were identical to Experiment 1.

Participants were randomly assigned to one of eight block sequences. These were all combinations of block orders with the restriction that both baseline conditions had to be presented either before or after the two moving conditions. The sequences were balanced across participants using a block-randomized design to ensure equal representation of the sequences.

In the sitting baseline, participants were seated with both feet in contact with the ground. This was done to preserve the correspondence with Experiment 1. In the standing baseline, participants stood in a posture consistent with the experimental conditions, holding their right foot elevated above the ground in front of them, at approximately the same position where it would collide with the moving ball (ca. 15 cm above ground; Figure 4). We introduced the standing baseline to control for any tactile modulation arising due to the postural demands or due to motor commands reducing sensitivity, therefore reducing the observed effects to the impact of the ball. We introduced breaks of 2-3 sec between trials of the standing baseline block, during which participants adopted a bipedal stance to rest.

For each experimental block (high and low speed), participants performed five practice trials to familiarize themselves with the ball speeds used: either 1500 mm/s or 2500 mm/s. These speeds correspond approximately to the 25th (1742 mm/s) and 75th (2579 mm/s) percentiles of contact speeds observed in Experiment 1 (see Results-Kinematics in Experiment 1). During the practice trials, participants were instructed to lift their right foot to the position in the air aligned with the edge of the occlude in medio-lateral dimension (Figure 4b), where the ball would make contact with it. Each trial started when the experimenter providing a verbal cue, which requested the participants to adopt the necessary leg posture. To achieve the desired ball speed, the experimenter had to lift the ball to a certain angle and then release it. Release height was variable and was adjusted on a trial-by-trial basis since the smoothness of ball release, exact foot placement, and exact rope lengths were slightly variable. Resulting speeds of the ball at the moment of *contact* can be seen in Figure 5b. Meanwhile, it was important that participants could not predict when the ball would start moving toward their foot and from what height, because otherwise they could establish predictions about the timing and intensity of collision, which might influence tactile sensitivity. To prevent the participant from engaging in predictive behavior based on knowing the exact timing of the ball’s collision, the experimenter and the ball’s starting position were hidden behind an occluder positioned to the participant’s left (Figure 4). An auditory signal was delivered to the experimenter via in-ear earplugs to assist the experimenter in positioning the ball at the specific height so that, upon release, it would travel from the participant’s left side and collide with the inner part of their foot at the desired target speed (1500 or 2500 mm/s). This initial angle was determined through prior piloting and was adjusted trial-by-trial as necessary to achieve the desired speed. As soon as the experimenter placed the ball at the suitable angle relative to the beam, a continuous auditory signal was delivered to the experimenter’s in-ear headphones to inform her that the ball is at the right place. This signal could not be heard by the participant. The experimenter then released the ball. Ball-foot contact was defined as the point at which the ball was within 1.5 cm of the previously established minimum ball-foot distance. At that moment, an intense vibrotactile probe (0.632 mm) was delivered to help participants understand the trial structure. These five practice trials of each experimental block also served to determine the ball’s position 235 ms prior to contact for each speed condition, using the averaged data from these trials. This was important for having valid estimates of the *before* time point during the experimental trials and being able to have the *before* condition as a first trial, as it had been done in earlier studies (Arikan et al., 2021; Gertz et al., 2017).

**Figure 5:**
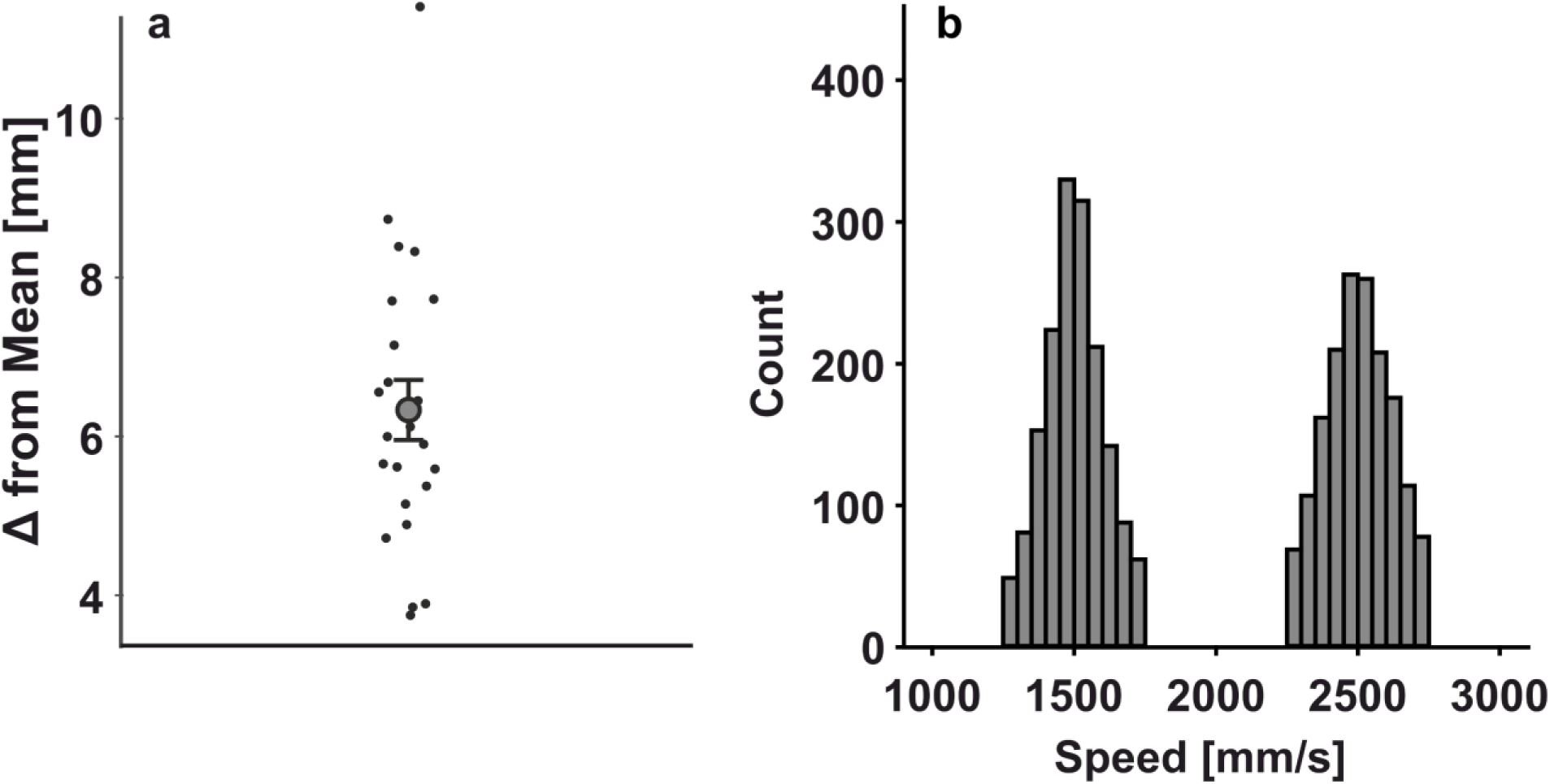
Paradigm analysis. (a) Maximal foot displacement to the mean position during the last 300 ms prior to ball contact averaged over all participants. Medians of single participants are depicted by single dots. The error bar shows the standard error while the circle indicates the mean. (b) Distribution of the ball speeds over all trials at the moment of contact.

The timing of 235 ms before contact was selected as it corresponded to the median stimulation time (236 ms) observed during the swing phase in Experiment 1. In each of the two experimental blocks, vibrotactile stimulation was randomly administered at one of three time points: *before* contact (-235 ms), at contact, or *after* contact (+250 ms). These time points approximated the phases of *swinging*, *contact*, and *after*-contact used in the first experiment.

As in the practice trials, during the experimental blocks, participants were instructed to hold their right foot elevated in front of them after receiving the experimenter’s signal and to wait for the ball to make contact with it. After each trial, participants were asked to indicate whether they perceived the stimulation. Stimulation intensity was decided using the same procedure as in the first experiment. Participants were encouraged to take a bipedal stance between trials to avoid fatigue.

With two ball speeds, three stimulation time points, and two baseline conditions, the experiment involved eight conditions, each managed by a separate QUEST algorithm. Each algorithm required 25 trials, resulting in a total of 200 trials per participant. Experiment 2 lasted approximately 30 min.

### Data analysis

As a first step in our analyses, we wanted to ensure that participants kept minimal foot movements around the time of ball contact. This was important to isolate possible effects of the collision between ball and foot from possible effects of moving the foot, as the former effects are related to masking whereas the latter can be related to sensorimotor processes, such as predictions stemming from motor commands associated with the active movement. To this end, we first calculated the maximal foot displacement during the 300 ms before ball contact (Figure 5a). We express this displacement relative to the foot mean position over the same period. Specifically, for each trial, we calculated the maximal 3D distance between the foot and the average foot position during the 300 ms before the ball contacted the foot. We then computed the median across all trials per participant. Additionally, we determined the actual timing of stimulation for each of the three time points of stimulation (*before*, at *contact*, *after*). This resulted in (mean ± standard deviation) -0.245 ± 0.199 ms, 0 ± 0 ms, and 0.252 ± 0.003 ms for the *before*, at *contact*, and *after* time points, respectively. Lastly, we verified whether the achieved and desired speeds at *contact* matched by differentiating the central ball position calculated from the 4 ball markers and computing the Euclidean norm across the three spatial dimensions (Figure 5b).

For the psychophysical analysis, we normalized the data to account for individual differences by subtracting the detection threshold obtained during the standing baseline from those estimated in each of the six experimental conditions. Hereby, we eliminated the influence that is associated with standing and elevating the foot. No participant was excluded from statistical testing.

### Statistical analysis

To assess whether the contact speed and the timing of stimulation influence tactile sensitivity on the static foot, we conducted a 2 (ball speed: slow, fast) × 3 (stimulation timing: *before, contact, after*) repeated-measures ANOVA. Additionally, to investigate whether higher stimulus intensity—expressed in this experiment as different ball impulses—led to greater tactile masking, we performed a one-sided paired-sample t-test on the normalized detection thresholds at the moment of *contact*. The other two time points were not analyzed this way, as they fall outside the temporal window where tactile masking has previously been reported (Abramsky et al., 1971; Kirman, 1984). To test if this is true in our dataset we conducted 2 frequentist and Bayesian paired t-tests between the 2 levels of ball speed intensities that could cause masking. If there was no effect of masking we would assume a Bayes Factor (BF_0+_) null versus alternative hypothesis above 3, following common recommendations (Andraszewicz et al., 2015) and p values exceeding the conventional threshold of 0.05. We also calculated a paired samples t-test to confirm and replicate an earlier finding that standing itself caused suppression compared to sitting (Wachsmann et al., 2025b).

### Results Experiment 2

The 2×3 repeated measure ANOVA confirmed the hypothesized main effect of time (*F*_2,44_ = 63.455, *p* < 0.001, *η^2^* = 0.743; Figure 6) and an interaction of time and masking stimulus intensity (*F*_2,22_ = 4.139, *p* = 0.023, *η^2^* = 0.158) that arose because suppression at ball *contact* was higher when the ball speed was higher than lower (*t*_22_ = 1.909, *p* = 0.035, *η^2^* = 0.038) while no differences were observed in the other two time points (both p > 0.33 and BF_0+_ > 3.152). The main effect of masking stimulus intensity (ball speed) did not reach significance (*F*_2,44_ = 2.406, *p* = 0.135 *η^2^* = 0.099). Detection thresholds at baseline were significantly higher in standing than sitting, as expected (*t*_22_ = 2.672, *p* = 0.007, *η^2^* = 0.072).

**Figure 6:**
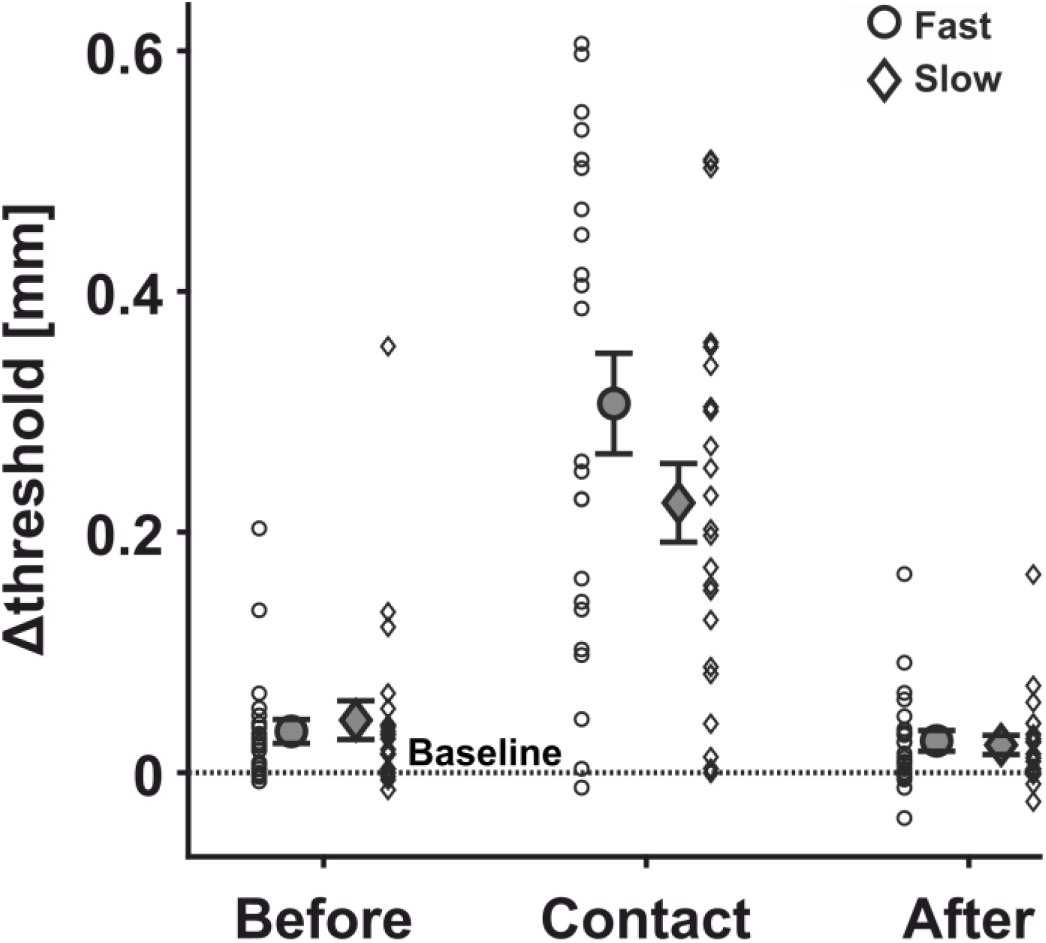
Psychophysical results. Detection thresholds during experimental trials normalized to the standing baseline (Δthreshold) at the three time points of stimulation relative to ball contact. Single subject data are depicted as small symbols (circle for fast and diamonds for slow ball velocities) while big symbols with error bars represent the means and standard errors. The dashed line shows baseline performance.

## Discussion Experiment 2

In the second experiment, we aimed to test whether tactile processing in the lower limbs is influenced by peripheral processes, such as tactile masking, which might explain the pronounced and variable suppression at the time of ball *contact*, and possibly adjacent time points, in Experiment 1. To do so, participants held their static foot in the air without visual information about a ball that approached their foot with two different speeds. We again applied tactile stimuli at three different time points.

As expected (Abramsky et al., 1971), we observed a clear increase in tactile detection thresholds at the time of ball *contact* compared to the time points *before* and *after* contact. This suggests that tactile sensitivity at *contact* is strongly influenced by masking. We did not find a main effect of masking stimulus intensity, which is unsurprising given that the *before* and *after* time points were outside the typical temporal ranges for forward and backward masking (Abramsky et al., 1971; Kirman, 1984). We could statistically confirm that the *before* and *after* time points were not due to ball contact-dependent masking, since the impulses did not influence tactile sensitivity. More importantly, we found a significant interaction between time point and ball speed, which is driven by suppression being larger when the ball contacted the foot with higher than lower speeds. This interaction highlights the fact that the influence of masking stimulus intensity on tactile sensitivity is most pronounced during ball *contact*, underscoring the temporal specificity of the masking phenomenon.

These findings indicate that tactile masking may have contributed to the pronounced suppression at the moment of ball *contact* observed in Experiment 1. However, more factors are likely involved in that finding, such as the precise prediction about when and how quickly the kicking foot would hit the ball. Indeed, more reliable predictions about the sensory consequences of one’s own action lead to stronger suppression (Blakemore et al., 1999; Fuehrer et al., 2022). Tactile masking at the foot during collisions with a ball is modulated by the masking stimulus intensity, and this occurs within a similar time window to that reported in the upper limbs (Craig, 1974; Fraser & Fiehler, 2018; Kirman, 1984; Schmid, 1961). We speculate that masking generalizes throughout the leg since reduction in tactile sensitivity also appears at the calf when postural sways are larger (Wachsmann et al., 2025b).

## Summary

In summary, we demonstrate that tactile sensitivity in the lower limbs is dynamically modulated during complex, everyday movements and that this modulation resembles those observed in upper limb goaldirected movements. In Experiment 1, we identified a distinct pattern in the kicking leg: tactile sensitivity increased from movement *onset* to the *swing* phase—when guiding demands are highest—followed by pronounced suppression at ball *contact* and a subsequent recovery of sensitivity. In contrast, tactile sensitivity was suppressed in the balancing foot during the kicking’s leg *swing* phase, with recovery thereafter. Notably, this suppression did not align with the peak foot speed, which occurs around movement *onset* of the kicking leg, suggesting that postural demands rather than movement speed primarily drive the modulation (Cybulska-Klosowicz et al., 2011).

Across both experiments, tactile suppression at the moment of ball *contact* scaled with contact impulse. Experiment 2 confirmed that this effect occurs within a narrow temporal window, as tactile sensitivity during the *before* and *after* time points was not substantially affected. This strengthens our confidence in the observed magnitudes of modulation effects of the first experiment. Together, two tactile sensitivity modulation patterns were observed in the different feet, suggesting that tactile modulation is governed by two main mechanisms: tactile sensitivity can be upweighted at moments when online feedback from the probed limb is important for the action (Colino & Binsted, 2016; Voudouris et al., 2019; Voudouris & Fiehler, 2022), while postural demands (Wachsmann et al., 2025b) and masking (Abramsky et al., 1971; Kirman, 1984) reduce it.

## Lead contact

Further information and requests for resources should be directed to and will be fulfilled by the lead contact, Fabian Dominik Wachsmann

## Acknowledgments

This work was supported by the Collaborative Research Center SFB/TRR 135, project A4, under grant agreement 222641018 funded by the German Research Foundation (Deutsche Forschungsgemeinschaft, DFG), by the German Research Foundation under Germany’s Excellence Strategy (EXC 3066/1 “The Adaptive Mind”, Project No. 533717223) and by the German Research Foundation (project VO 2542/1-1). We would like to thank Fabian Krause and Clara Lang for their assistance in data collection.

## Data availability

Behavioral and psychophysical data are publicly available after acceptance at osf.io/bvtdk. For further information or data requests please correspond to Fabian Dominik Wachsmann.

## Author contributions

Conceptualization: F.W., K.F., D.V.

Data curation: F.W.

Formal analysis: F.W.

Funding acquisition: K.F., D.V.

Investigation: F.W.

Methodology: F.W., K.F., D.V.

Project administration: F.W., K.F., D.V.

Resources: F.W., K.F., D.V.

Software: F.W.

Supervision: K.F., D.V.

Validation: F.W., K.F., D.V.

Visualization: F.W.

Roles/Writing original draft: F.W.

and Writing review & editing: F.W., K.F., D.V.

## Appendix

**Appendix 1:**
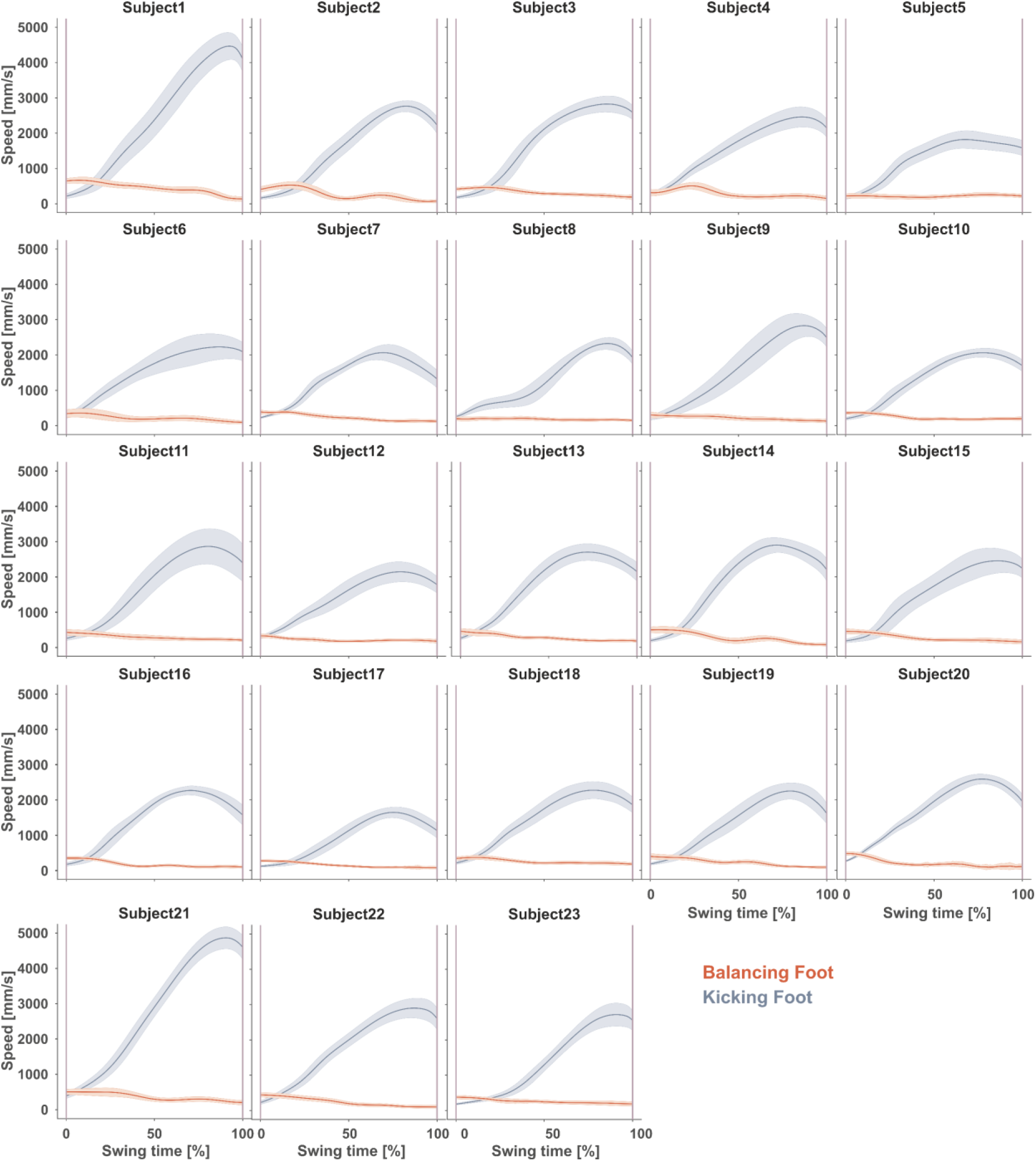
Single subject speed profiles. The figure shows the time normalized single subject time-courses of the swinging phase for the kicking (blue) and the balancing (red) leg for all participants, with standard errors as shaded error bars. The first and second purple vertical lines indicate movement onset and moment of ball contact, respectively. Please note that the speed of the balancing leg is decreasing as it reache d maximum around the kicking foot movement onset.

**Appendix 2:**
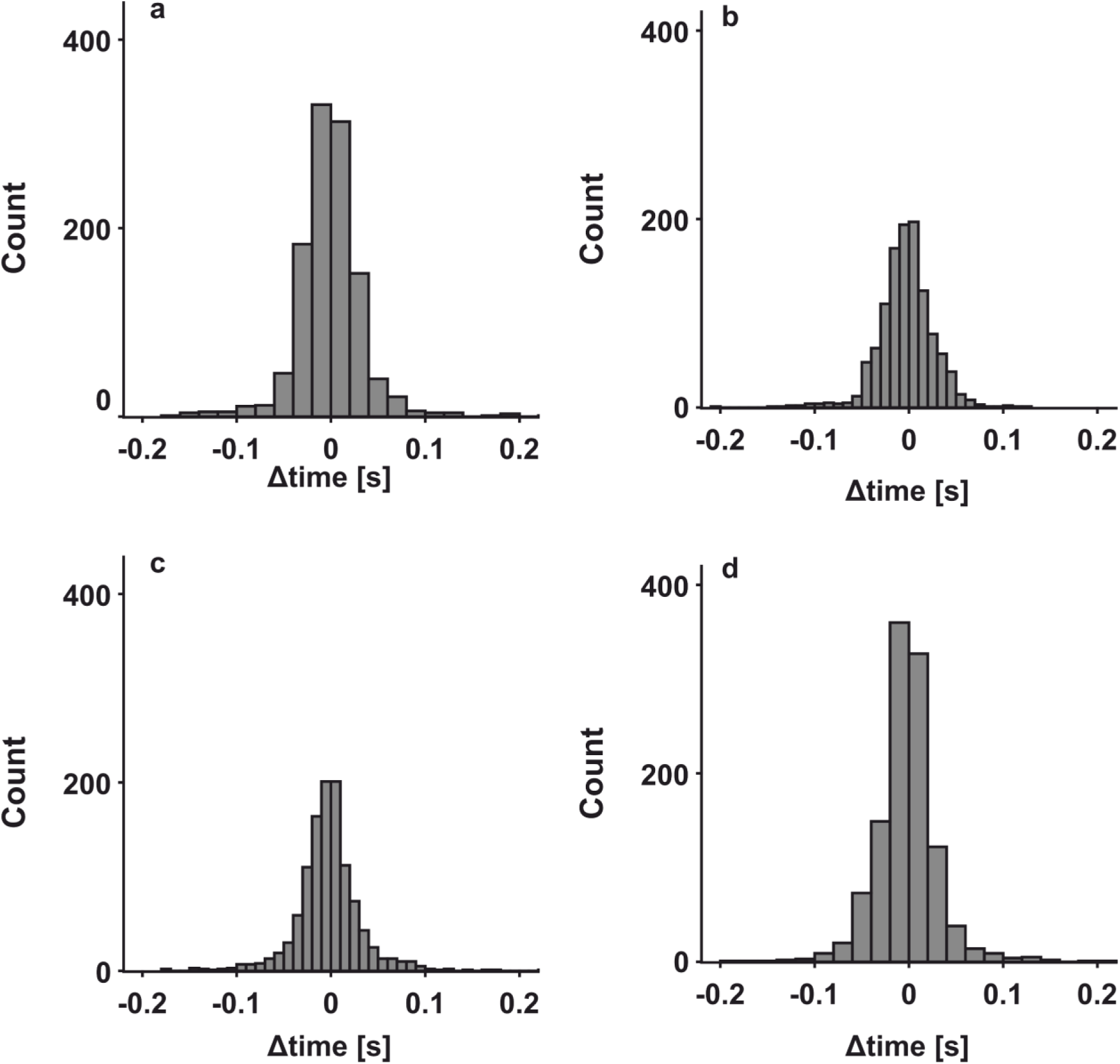
Differences between intended and actual stimulation. Histograms of the time differences between the intended and the actual vibrotactile stimulation for all trials, separately for the four stimulation time points (a) onset, (b) swing, (c) ball contact, and (d) after-contact.

**Appendix 3:**
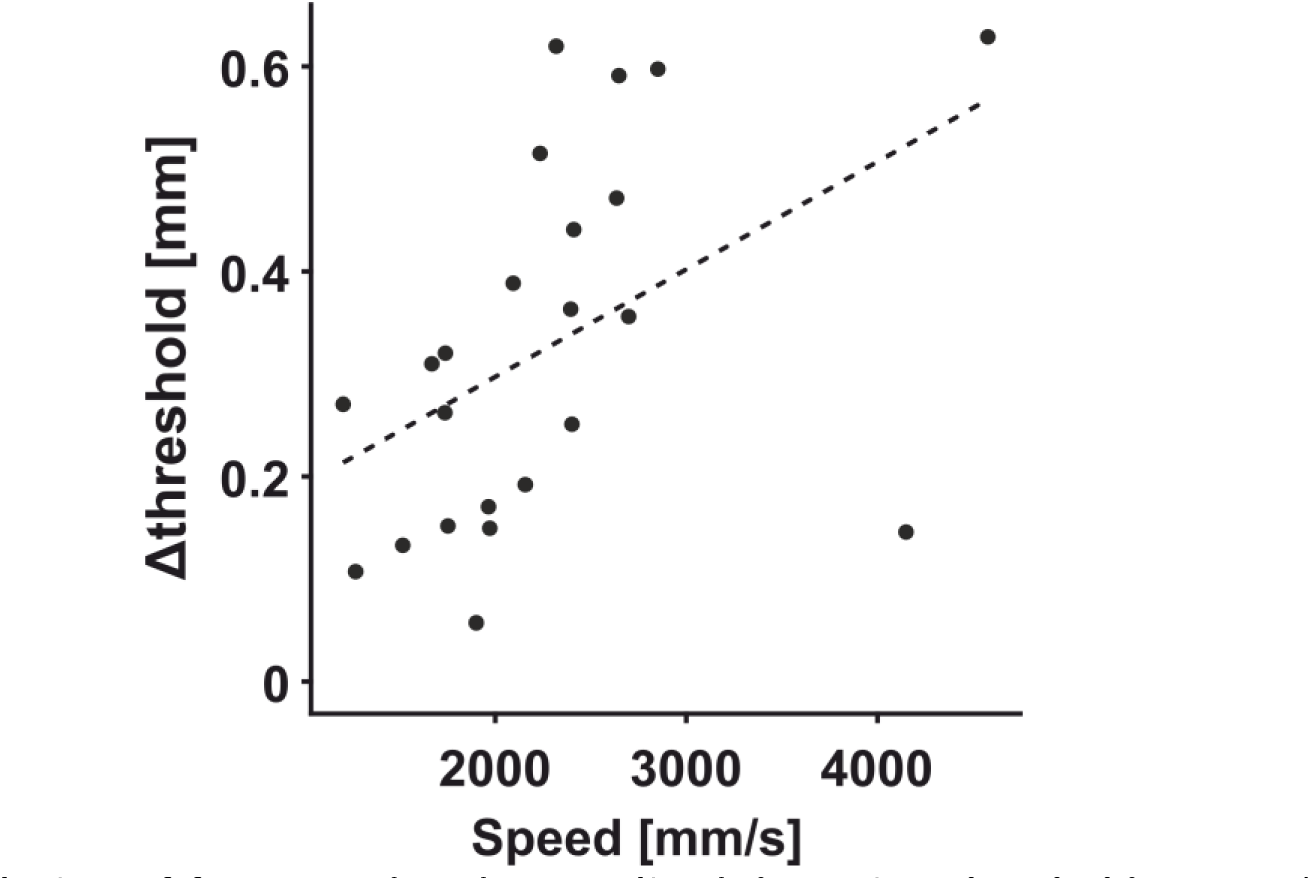
Correlation of foot speed and normalized detection threshold. Normalized detection thresholds as a function of the kicking leg’ s speed at the moment of ball contact. The strength of suppression is expressed as the amplitude of the vibration stimulus difference from baseline. Dots represent single subject data, while the dashed line indicates a first order polynomial fitted to the data.

## References

1. Abramsky, O., Carmon, A., & Benton, A. L. (1971). Masking of and by tactile pressure stimuli. Perception & Psychophysics, 10(5), 353–355. 10.3758/BF03207457

2. Andraszewicz, S., Scheibehenne, B., Rieskamp, J., Grasman, R., Verhagen, J., & Wagenmakers, E.-J. (2015). An Introduction to Bayesian Hypothesis Testing for Management Research. Journal of Management, 41(2), 521–543. 10.1177/0149206314560412

3. Arikan, B. E., Voudouris, D., Straube, B., & Fiehler, K. (2024). Distinct role of central predictive mechanisms in tactile suppression. IScience, 27(8), 110582. 10.1016/j.isci.2024.110582

4. Arikan, B. E., Voudouris, D., Voudouri-Gertz, H., Sommer, J., & Fiehler, K. (2021). Reach-relevant somatosensory signals modulate activity in the tactile suppression network. NeuroImage, 236, 118000. 10.1016/j.neuroimage.2021.118000

5. Blakemore, S., Frith, C. D., & Wolpert, D. M. (1999). Spatio-temporal prediction modulates the perception of self-produced stimuli. Journal of Cognitive Neuroscience, 11(5), 551–559. 10.1162/089892999563607

6. Blakemore, S., Wolpert, D., & Frith, C. (2000). Why can’t you tickle yourself?: NeuroReport, 11(11), R11–R16. 10.1097/00001756-200008030-00002

7. Buckingham, G., Carey, D. P., Colino, F. L., deGrosbois, J., & Binsted, G. (2010). Gating of vibrotactile detection during visually guided bimanual reaches. Experimental Brain Research, 201(3), 411–419. 10.1007/s00221-009-2050-8

8. Chapman, C. E., Bushnell, M. C., Miron, D., Duncan, G. H., & Lund, J. P. (1987). Sensory perception during movement in man. Experimental Brain Research, 68(3), 516–524. 10.1007/BF00249795

9. Chew-Bullock, T. S.-Y., Anderson, D. I., Hamel, K. A., Gorelick, M. L., Wallace, S. A., & Sidaway, B. (2012). Kicking performance in relation to balance ability over the support leg. Human Movement Science, 31(6), 1615–1623. 10.1016/j.humov.2012.07.001

10. Colino, F. L., & Binsted, G. (2016). Time Course of Tactile Gating in a Reach-to-Grasp and Lift Task. Journal of Motor Behavior, 48(5), 390–400. 10.1080/00222895.2015.1113917

11. Craig, J. C. (1974). Vibrotactile difference thresholds for intensity and the effect of a masking stimulus. Perception & Psychophysics, 15(1), 123–127. 10.3758/BF03205839

12. Cybulska-Klosowicz, A., Meftah, E.-M., Raby, M., Lemieux, M.-L., & Chapman, C. E. (2011). A critical speed for gating of tactile detection during voluntary movement. Experimental Brain Research, 210(2), 291–301. 10.1007/s00221-011-2632-0

13. Elliott, D., Lyons, J., Hayes, S. J., Burkitt, J. J., Roberts, J. W., Grierson, L. E. M., Hansen, S., & Bennett, S. J. (2017). The multiple process model of goal-directed reaching revisited. Neuroscience & Biobehavioral Reviews, 72, 95–110. 10.1016/j.neubiorev.2016.11.016

14. Fraser, L. E., & Fiehler, K. (2018). Predicted reach consequences drive time course of tactile suppression. Behavioural Brain Research, 350, 54–64. 10.1016/j.bbr.2018.05.010

15. Fuehrer, E., Voudouris, D., Lezkan, A., Drewing, K., & Fiehler, K. (2022). Tactile suppression stems from specific sensorimotor predictions. Proceedings of the National Academy of Sciences, 119(20), e2118445119. 10.1073/pnas.2118445119

16. Gertz, H., Voudouris, D., & Fiehler, K. (2017). Reach-relevant somatosensory signals modulate tactile suppression. Journal of Neurophysiology, 117(6), 2262–2268. 10.1152/jn.00052.2017

17. Hennig, E. M., & Sterzing, T. (2009). Sensitivity Mapping of the Human Foot: Thresholds at 30 Skin Locations. Foot & Ankle International, 30(10), 986–991. 10.3113/FAI.2009.0986

18. Kellis, E., & Katis, A. (2007). Biomechanical characteristics and determinants of instep soccer kick. Journal of Sports Science & Medicine, 6(2), 154–165.

19. Kilteni, K., & Ehrsson, H. H. (2022). Predictive attenuation of touch and tactile gating are distinct perceptual phenomena. IScience, 25(4), 104077. 10.1016/j.isci.2022.104077

20. Kirman, J. H. (1984). Forward and Backward Tactile Recognition Masking. The Journal of General Psychology, 111(1), 83–99. 10.1080/00221309.1984.9921100

21. Kozłowska, A. (1998). Studying tactile sensitivity – population approach. Anthropological Review, 61, 3–30. 10.18778/1898-6773.61.01

22. Menz, H. B., Lord, S. R., & Fitzpatrick, R. C. (2006). A tactile stimulus applied to the leg improves postural stability in young, old and neuropathic subjects. Neuroscience Letters, 406(1–2), 23–26. 10.1016/j.neulet.2006.07.014

23. Mouchnino, L., Fontan, A., Tandonnet, C., Perrier, J., Saradjian, A. H., Blouin, J., & Simoneau, M. (2015). Facilitation of cutaneous inputs during the planning phase of gait initiation. Journal of Neurophysiology, 114(1), 301–308. 10.1152/jn.00668.2014

24. Pearcey, G. E. P., & Zehr, E. P. (2019). We Are Upright-Walking Cats: Human Limbs as Sensory Antennae During Locomotion. Physiology, 34(5), 354–364. 10.1152/physiol.00008.2019

25. Peterka, R. J. (2002). Sensorimotor Integration in Human Postural Control. Journal of Neurophysiology, 88(3), 1097–1118. 10.1152/jn.2002.88.3.1097

26. Schmid, E. (1961). Temporal aspects of cutaneous interaction with two-point electrical stimulation. Journal of Experimental Psychology, 61(5), 400–409. 10.1037/h0043334

27. Shergill, S. S., Bays, P. M., Frith, C. D., & Wolpert, D. M. (2003). Two Eyes for an Eye: The Neuroscience of Force Escalation. Science, 301(5630), 187–187. 10.1126/science.1085327

28. Voudouris, D., Broda, M. D., & Fiehler, K. (2019). Anticipatory grasping control modulates somatosensory perception. Journal of Vision, 19(5), 1–10. 10.1167/19.5.4

29. Voudouris, D., & Fiehler, K. (2017). Enhancement and suppression of tactile signals during reaching. Journal of Experimental Psychology: Human Perception and Performance, 43(6), 1238–1248. 10.1037/xhp0000373

30. Voudouris, D., & Fiehler, K. (2021). Dynamic temporal modulation of somatosensory processing during reaching. Scientific Reports, 11, 1928. 10.1038/s41598-021-81156-0

31. Voudouris, D., & Fiehler, K. (2022). The role of grasping demands on tactile suppression. Human Movement Science, 83, 102957. 10.1016/j.humov.2022.102957

32. Wachsmann, F. D., Fiehler, K., & Voudouris, D. (2025a). Temporal modulation of tactile perception during balance control. Scientific Reports, 15(1), 17380. 10.1038/s41598-025-99006-8

33. Wachsmann, F. D., Fiehler, K., & Voudouris, D. (2025b). Postural demands modulate tactile perception in the lower limb in young and older adults. Scientific Reports, 15(1), 20221. 10.1038/s41598-025-06736-w

34. Watson, A. B., & Pelli, D. G. (1983). Quest: A Bayesian adaptive psychometric method. Perception & Psychophysics, 33(2), 113–120. 10.3758/BF03202828

35. World Medical Association Declaration of Helsinki: Ethical Principles for Medical Research Involving Human Subjects. (2013). JAMA, 310(20), 2191. 10.1001/jama.2013.281053

